# Mechanism of LLPS of SARS-CoV-2 N protein

**DOI:** 10.1101/2022.12.21.521431

**Authors:** Mei Dang, Tongyang Li, Jianxing Song

**Affiliations:** Department of Biological Sciences, Faculty of Science; National University of Singapore; 10 Kent Ridge Crescent, Singapore 119260

**Keywords:** ATP: Nucleic acid, SARS-CoV-2, Nucleocapsid (N) protein, Liquid-liquid phase separation (LLPS), NMR spectroscopy

## Abstract

SARS-CoV-2 nucleocapsid (N) protein with low mutation rate is the only structural protein not only functioning to package viral genomic RNA, but also manipulating the host-cell machineries, thus representing a key target for drug development. Recent discovery of its liquid-liquid phase separation (LLPS) not only sheds light on previously-unknown mechanisms underlying the host-SARS-CoV-2 interaction and viral life cycle, but most importantly opens up a new direction for developing anti-SARS-CoV-2 strategies/drugs. However, so far the high-resolution mechanism of LLPS of N protein still remains unknown because it is not amenable for high-resolution biophysical investigations. Here we systematically dissected N protein into differential combinations of domains followed by DIC and NMR characterization. We successfully identified N (1-249), which not only gives high-quality NMR spectra, but phase separates as the full-length N protein. The results together decode for the first time: 1) nucleic acid modulates LLPS by dynamic but specific interactions multivalently over both folded NTD/CTD and Arg/Lys residues within IDRs. 2) ATP, mysteriously with concentrations >mM in all living cells but absent in viruses, not only specifically binds NTD/CTD, but also Arg residues within IDRs with Kd of 2.8 mM. 3) ATP dissolves LLPS by competitively displacing nucleic acid from binding the protein. Therefore, ATP and nucleic acid interplay in modulating LLPS by specific competitions for binding over the highly overlapped binding sites. Our study deciphers the mechanism of LLPS of N protein, which is targetable by small molecules. ATP is not only emerging as a cellular factor controlling the host-SARS-CoV-2 interaction, but also provides a lead for developing anti-SARS-CoV-2 drugs efficient for different variants of SARS-CoV-2. Fundamentally, our results imply that the mechanisms of LLPS of IDR-containing proteins mediated by ATP and nucleic acids appear to be highly conserved from human to virus.

## Introduction

Severe Acute Respiratory Syndrome Coronavirus 2 (SARS-CoV-2) is a member of a large family of coronaviruses with ∼ 30 kb genomic RNA (gRNA) packaged by nucleocapsid (N) protein in a membrane-enveloped virion, which caused the ongoing pandemic with >300 millions of infections and >5.47 millions of deaths (1). SARS-CoV-2 has four structural proteins: namely the spike (S) protein that recognizes the host-cell receptors angiotensin converting enzyme-2 (ACE2), membrane-associated envelope (E), membrane (M) proteins and nucleocapsid N protein. In order to determine their potential for development of vaccine and drug, there is an urgent need to understand structures and functions as well as roles in the viral life cycle of SARS-CoV-2 proteins beyond the spike protein which is currently used for vaccine.

N protein is the only structural protein which not only functions to package gRNA, but is also responsible for suppressing the immune system and manipulating the cellular machineries of the host cell to enhance the viral infection and replication. For example, N protein has been currently identified to play critical roles in hijacking the host cell machineries for RNA replication and transcription, nucleocapsid assembly, virion assembly and virion package. Furthermore, N protein provides the connection between viral E, M proteins and gRNA within virion. In particular, it has high immunogenicity (4) and a low rate of mutation (Fig. S1A), with 91% identity to that of SARS-CoV-1, which is much more conserved than the S protein. Consequently, N protein represents a key candidate for future drug and vaccine development (2-16).

Most strikingly, liquid-liquid phase separation (LLPS), the emerging principle for commonly organizing the membrane-less organelles (MLOs) critical for cellular physiology and pathology (17-19), has been very recently identified as the key mechanism underlying the diverse functions of SARS-CoV-2 N protein (10-16). Noticeably, most, if not all, functions of N proteins including LLPS appear to be dependent on its binding to various viral/host-cell nucleic acids including single- and double-stranded RNA/DNA of diverse sequences. For example, SARS-CoV-2 N protein has been previously shown to phase separate upon introducing various nucleic acids of specific and non-specific sequences. Moreover, although the detailed mechanism still remains poorly understood, the final package of the RNA genome requires the complex but precise interaction between N protein and gRNA, which should be extremely challenging for SARS-CoV-2 with such a large RNA genome (∼30 kb). In this context, any small molecules capable of modulating the interaction of N protein with nucleic acids is anticipated to significantly intervene in key steps of the viral life cycle, some of which might be further developed into anti-SARS-CoV-2 drugs.

ATP, the universal “biological fuel” for all living cells, has cellular concentrations from 2 to 16 mM depending on the types of cells, which are much higher than required for its classic functions. By contrast, viruses lack the ability to generate ATP (20). Emerging results indicate that ATP appears to have a novel category of energy-independently functions at mM, which include the modulation of LLPS and specific binding to a list of the nucleic-acid-binding domains of diverse folds at mM (19,21-27). Very recently we found that ATP is in fact capable of specifically binding to pockets on its NTD (10) and CTD (27) with the dissociation constant (Kd) of 3.3 ± 0.4 and 1.49 ± 0.28 mM as well as dissolving LLPS of SARS-CoV-2 N protein but with the mechanism unknown. These results imply that at mM ATP might not only have functions to control protein hemostasis in living cells, but also act as a key host-cell factor controlling the interaction of the host cell with viruses such as SARS-CoV-2 which are unable to generate ATP.

SARS-CoV-2 N protein is a 419-residue multidomain protein (Fig. 1A) composed of the folded N-terminal domain (NTD) and C-terminal domain (CTD), as well as three long intrinsically-disordered regions (IDRs) containing a number of Arg/Lys residues (Fig. 1B and SFig. 1A). Previous studies have established that its NTD is a nucleic-acid-binding domain (RBD) functioning to interact with various RNA and DNA of specific and non-specific sequences (6-8), while CTD was previously established to dimerize/oligomerize to form high-order structures (8,9). Nevertheless, very recently, we found that its CTD is also a cryptic domain for binding ATP and nucleic acids (27). Very importantly, the sequences of N protein are highly conserved in different variants including Delta and Omicron not only over the folded NTD and CTD, but also over three IDRs (Fig. 1A). This observation implies that IDRs also have critical functional roles which remain to be discovered.

**Figure 1.**
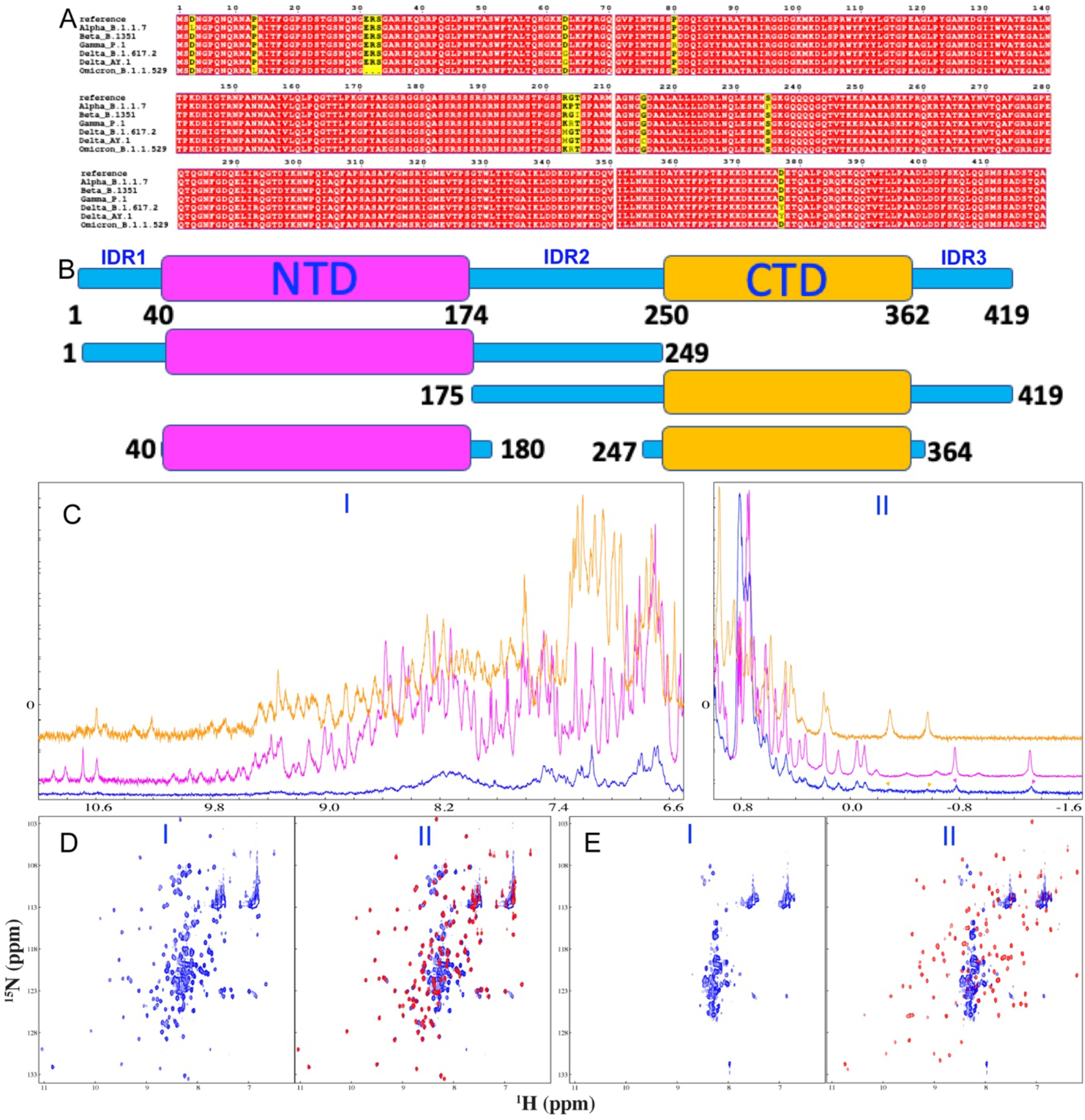
NMR characterization of differentially-dissected domains of SARS-CoV-2 N protein. (A) Sequence alignment of N protein of Variants of Concern (VOCs) of SARS-CoV-2 according to WHO: https://www.who.int/en/activities/tracking-SARS-CoV-2-variants. (B) Differentially-dissected domains of SARS-CoV-2 N protein used in this study. (C) One dimensional NMR proton spectra of the full-length N protein (blue), NTD (purple) and CTD (light brown) for the amide proton (I) and side-chain (II) regions. Arrows are used to indicate very up-field signature NMR signals of methyl groups of NTD (purple) and CTD (light brown) in 1D NMR spectrum of the full-length N protein. (D) HSQC spectrum of N (1-249) (I) and superimposition of HSQC spectra of N( 1-249) (blue) and NTD (44-180) (red) (II). (E) HSQC spectrum of N (175-419) (I) and superimposition of HSQC spectra of N (175-419) (blue) and CTD (247-364) (red) (II).

Despite the criticality of understanding LLPS of N protein for developing therapeutic strategies/molecules to fight the pandemic, the high-resolution mechanisms for LLPS of N protein still remain unknown most likely due to the challenge in characterizing the association-prone N protein by the high-resolution biophysical methods particularly by NMR spectroscopy, the only one available so far to experimentally obtain the residue-specific knowledge of LLPS (12,21-28). Here, to overcome the challenge, we first dissected N protein into the differential combinations of domains followed by NMR and DIC characterization. Consequently we have successfully identified N (1-249) composed of NTD and IDRs, which not only gives high-quality NMR spectra, but also phase separates as the full-length N protein. Subsequently we carried out extensive NMR and DIC studies on its LLPS induced by S2m, a 32-mer specific stem-loop II nucleic acid motif (Fig. S1B) derived from SARS-CoV-2 gRNA (27,29) as well as modulated by ATP. The obtained results, for the first time, to the best of our knowledge, decode these novel findings: 1) the structures of NTD and CTD are highly similar in both isolated domains and full-length N protein. However, the presence of IDRs together with CTD triggers further dynamic oligomerization or/and conformational exchanges, thus making NMR signals of the full-length N protein too broad to be suitable for high-resolution NMR investigations. 2) Nevertheless, N (1-249) is not only amenable for high-resolution NMR studies, but also phase separate upon induction by S2m and modulation by ATP in the same manner as the full-length N protein. 3) ATP specifically binds the nucleic-acid-binding pocket of the folded NTD as well as Arg residues within IDRs in the context of N (1-249) with the average Kd values of 2.0 ± 0.2 and 2.8 ± 0.2 mM respectively. 4) Residues-specific NMR results reveals that S2m induces and subsequently dissolves LLPS by multivalent and specific binding to both folded NTD and Arg/Lys residues within IDRs. 5) NMR visualization unambiguously reveals that ATP dissolves S2m-induced LLPS by competitively displacing S2m from being bound with the protein. These results indicate that LLPS of SARS-CoV-2 N protein is mainly driven by dynamic but specific interactions with nucleic acid multivalently over both NTD/CTD and IDRs. ATP dissolves LLPS of N protein also by specifically competing with nucleic acid for binding the highly overlapped sites on NTD/CTD and IDRs. In light of the mechanism of LLPS of human FUS protein we previously revealed (21-23), our present study implies that the mechanisms of LLPS induced by nucleic acids for the proteins composed of the folded nucleic-acid-binding domains and Arg/Lys residues within IDRs are highly conserved from human to virus. Intriguingly, although SARS-CoV-2 cannot generate ATP, ATP is not only emerging as a key cellular factor controlling the host-SARS-CoV-2 interaction, but also provides the rationale/lead for further development of anti-SARS-CoV-2 drugs by targeting the ATP-binding sites of N protein.

## Results

### Dissection of N protein and their NMR characterization

In this study, we aimed to gain insights into the high-resolution mechanisms for LLPS of SARS-CoV-2 N protein that is induced by nucleic acid and modulated by ATP with NMR spectroscopy. Initially we attempted to carry out the study on the full-length N protein. Unfortunately, even after an extensive screening of protein concentrations and buffer conditions, the NMR resonance signals of the full-length N protein were very broad as exemplified by its one-dimensional NMR proton spectra (Fig. 1C), and consequently the signals of its amide protons were too broad to be detected in HSQC spectrum even without phase separation, completely consistent with a just published NMR study (7).

To identify the domains suitable for high-resolution NMR studies, we first dissected the full-length N protein into two large fragments consisting of differential domains, namely N (1-249) with IDR1, NTD and IDR2 as well as N (175-419) with IDR2, CTD and IDR3, together with the isolated NTD and CTD (Fig. 1B). The isolated NTD and CTD both have well-dispersed HSQC spectra even with peaks highly superimposable to those of the isolated NTD and CTD previously collected in slightly different buffers (6,30,31). In particular, in their one-dimensional proton NMR spectra (Fig. 1C), NTD has two very up-field signature signals respectively at −0.78 and −1.37 ppm while CTD has two at −0.25 and −0.58 ppm, which are all from the methyl groups with the close contact to the aromatic rings only observed in the well-folded proteins. Strikingly N (1-249) has a well-dispersed HSQC spectrum (I of Fig. 1D) and in particular, the HSQC peaks of its NTD residues are highly superimposable to those of the isolated NTD residues (II of Fig. 1D), indicating that the structures of NTD are highly similar in both isolated NTD and N (1-249).

By contrast, N (175-419) has a narrowly-dispersed HSQC spectrum (I of Fig. 1E) in which the well-dispersed peaks of the isolated CTD were completely undetected. Nevertheless, a close examination showed that the signature peaks of CTD could be still observed in the 1D spectra of the full-length (II of Fig. 1C) as well as (175-419) (Fig. S2), indicating that CTD is also similarly folded in the isolated domain, N (175-414) and the full-length N proteins. To confirm this, we further dissected N (175-419) into N (175-364) and N (247-419). As shown by their 1D spectra (Fig. S3A), N (175-464) and N (247-419) all have the signature peaks respectively at −0.25 and −0.58 ppm characteristic of the folded CTD which are less broad than those of N (175-419). Consistent with this, some well-dispersed HSQC peaks became detectable for N (175-464) and N (247-419), which are largely superimposable to those of the isolated CTD (Fig. S3B and S3C).

As such, most likely due to the self-association/oligomerization or/and μs-ms conformational dynamics provoked by the presence of IDRs and CTD, their NMR signals became broadened to different degrees for N (175-354), N (247-419), N (175-419) and full-length N proteins. Intriguingly, even in N (1-249) while most of HSQC peaks of the non-proline NTD and IDR1 residues could be detected, HSQC peaks of several segments of IDR2 were too broad to be detected, which include Ser184-Ser197, Leu221-Leu222, and Leu224-G236, thus implying that these segments are involved in μs-ms exchanges/dynamics as recently revealed by NMR backbone dynamics (7,30). Strikingly, these segments contain several hydrophobic residues Leu and particularly are rich in Ser residues (Fig. 1A) characteristic of the prion-like domains, which, as previously found, are prone to the self-association to form amyloid fibrils (32). Similarly, a very recent study also showed that CTD with IDRs together led to the disappearance of NMR signals of the full-length N protein although NTD and CTD were shown to have no significant interaction with each other (7). Nevertheless, the dissection study led to the successful identification of N (1-249) to be the largest fragment of N protein amenable for high-resolution NMR studies, which is not only composed of the folded NTD functioning to bind nucleic acids but also IDR1 and IDR2 essential for LLPS.

### ATP specifically binds residues of both folded NTD and IDRs

Previously we have found that ATP specifically binds the nucleic-acid-binding pocket of the isolated NTD with HSQC peaks of 11 residues significantly perturbed (10). Here we titrated N (1-249) at the same protein concentration (100 μM) and in the same buffer with ATP at 0.5, 1.0, 2.0, 4.0, 6.0, 10.0 mM respectively. Briefly, at ATP concentrations below 1 mM, only very minor shift of HSQC peaks was observed. However, upon further increasing ATP concentration, a small set of HSQC peaks underwent significant shift as illustrated by I of Fig. 2. Strikingly, comparison with the spectrum of the isolated NTD in the presence of ATP at 10 mM revealed that even in the context of N (1-249), the shift pattern of HSQC peaks of the NTD residues are highly similar to that of the isolated NTD (II of Fig. 2). Noticeably, on the other hand, in addition to the NTD residues, several peaks of the residues within IDRs unique for N (1-249) also underwent significant shift (III of Fig. 2).

**Figure 2.**
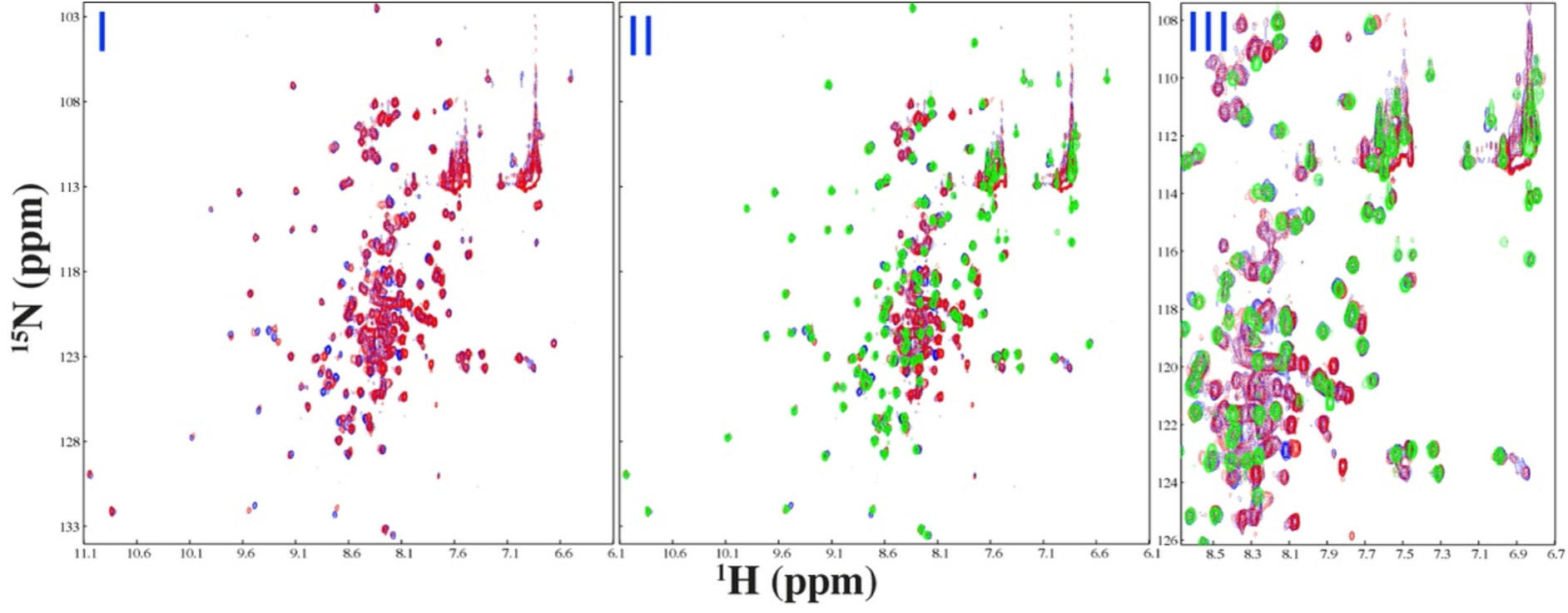
ATP specifically binds both NTD and IDR residues of N (1-249). (I) Superimposition of HSQC spectra of N (1-249) in the free state (blue) and in the presence of ATP at 10 mM (red). (II) Superimposition of HSQC spectra of N (1-249) in the free state (blue) and in the presence of ATP at 10 mM (red), as well as of NTD in the presence of ATP at 10 mM. (III) Zoom of HSQC spectra showing peaks of IDR residues of N (1-249) in the free state (blue) and in the presence of ATP at 10 mM (red), as well as of NTD in the presence of ATP at 10 mM (green).

Detailed analysis of HSQC spectra of N (1-249) in the absence and in the presence of ATP at different concentrations led to identification of significantly-perturbed residue of N (1-249). Indeed, the overall pattern of shifted HSQC peaks of the NTD residues are very similar in both isolated NTD and N (1-249) (Fig. 3A). Precisely, 10 NTD residues have significant shift including all those previously identified 11 residues in the isolated NTD except for Ala90, namely Asn48, Ser51, Leu56, Thr57, Arg89, Arg92, Ser105, Arg107, Ala155 and Tyr172. Comparison of the shift tracings of six representative residues showed that in the context of N (1-249) their shift tracings approached being saturated at the slightly lower ATP concentrations than for the isolated NTD (Fig. 3B).

**Figure 3.**
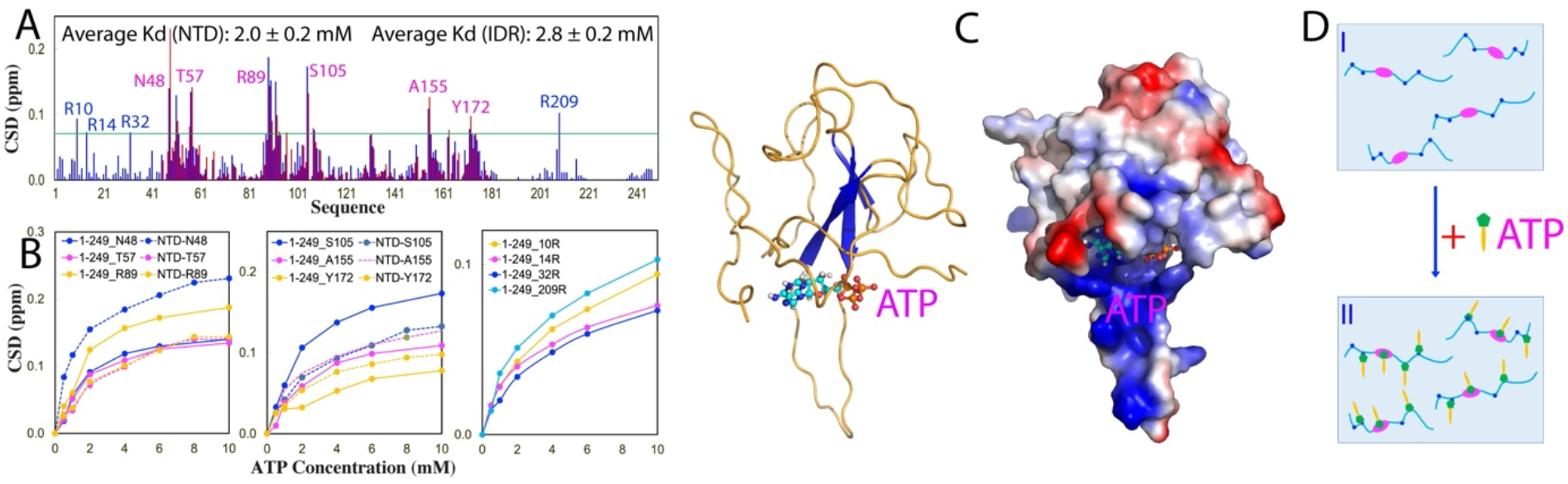
Residue-specific view of the ATP binding to NTD and IDR residues of N (1-249). (A) Residue-specific chemical shift difference (CSD) of N (1-249) (blue) and isolated NTD (red) between the free state and in the presence of ATP 10 mM. The significantly perturbed residues are defined as those with the CSD values at 10 mM > 0.072 (average value + one standard deviation) (cyan line). (B) Shift tracings of HSQC peaks of six representative NTD residue in the context of N (1-249) and isolated NTD, as well as 4 Arg residues within IDRs in the presence of ATP at different concentrations. (C) The docking structure of the ATP-NTD complex we previously constructed for the isolated NTD. (D) Schematic representation of N (1-249) in the free state (I) and in the presence of the exceeding amounts of ATP (II).

Indeed, the fitting of the shift tracings of 10 NTD residues in N (1-249) gave the average Kd of 2.0 ± 0.2 mM (Table S1), which is slightly smaller than that for the isolated NTD (3.3 ± 0.4 mM). This implies that ATP binds NTD in N (1-249) with the slightly higher affinity than the isolated NTD, which is similar to our previous observations that ATP binds the tandem-linked RRM domains with the slightly higher affinity than the isolated ones of TDP-43 and hnRNPA1 (33). Most strikingly, ATP also induced the significant shift of four Arg residues respectively within IDR1 and IDR2, namely Arg10, Arg14, Arg32 and Arg209 (Fig. 3A). Importantly, the fitting of their shift tracings (Fig. 3B) gave the average Kd of 2.8 ± 0.2 mM (Table S1), which is only slightly larger than that of the NTD residues in the same context of N (1-249).

The results together reveal that ATP is able to bind the nucleic-acid-binding pocket of the folded NTD in the context of N (1-249) with the complex structure very similar to what we previously constructed for the isolated NTD (Fig. 3C). Furthermore, here we decoded for the first time that ATP is not only able to bind the folded NTD, but also capable of binding Arg residues within IDRs with Kd of 2.8 mM (Fig. 3D), whose affinity is comparable to those for binding a list of the folded nucleic-acid-binding domains recently identified (22,23).

### ATP also binds the folded CTD in N (175-419)

Very recently, we shown that in addition to acting for dimerization/oligomerization, CTD in fact is a cryptic domain for binding ATP and S2m (27). In particular, CTD binds ATP with Kd of 1.49 ± 0.28 mM (27). Here we asked the question whether ATP can bind N (175-419). As evidenced by its 1D spectra in the presence of ATP at different concentrations (Fig. S4A), ATP did specifically induce the shift of one very up-field NMR peak at −0.25 but not the one at −0.58 ppm, exactly as we previously observed on the isolated CTD (27). This result clearly indicates that ATP is able to bind the folded CTD in the context of N (175-419). On the other hand, however, as N (175-419) have HSQC spectrum with many peaks undetectable likely due to μs-ms conformational dynamics or/and oligomerization, although ATP also triggered the broadening and disappearance of some HSQC peaks of IDRs (Fig. S4B-S4E), it is impossible to assign these peaks to the corresponding residues.

### S2m modulates LLPS of the full-length N and N (1-249) proteins

Very recently, SARS-CoV-2 N protein was shown to achieve its functionality through LLPS, which is driven by interacting with various nucleic acids including viral and host-cell RNA/DNA as well as non-specific sequences. For example, previously we found that A24, a 24-mer non-specific nucleic acid, was sufficient to achieve the biphasic modulation of LLPS of N protein: induction at low concentrations but dissolution at high concentrations (10). Here, we utilized S2m, a specific nucleic acid probe which was previously derived from gRNA to identify nucleic-acid-binding domains of SARS-CoVs (27,29) to assess its modulation of LLPS in parallel for the isolated NTD and CTD as well as the full-length N and N (1-249) proteins monitored by measuring the turbidity (absorption at 600 nm) and imaging with DIC microscopy, as we previously conducted on SARS-CoV-2 N protein (10), FUS (21) and TDP-43 (25).

We first titrated S2m into the isolated NTD and CTD with protein concentrations reaching up to 200 μM but found no phase separation. By contrast, as shown in Fig. 4, S2m could biphasically modulate LLPS of the full-length N protein: induction at low ratios and dissolution at high ratios. Briefly, the N protein sample showed no LLPS in the free state. However, phase separation was induced upon addition of S2m as evident by the increase of turbidity and DIC imaging. At 1:0.75 (N:S2m), the turbidity reached the highest of 1.92 (Fig. 4A) and many liquid droplets with the diameter of ∼1 μm were formed (Fig. 4B). However, further addition of S2m led to the reduction of turbidity and dissolution of the droplets. At 1:1.5 all liquid droplets were completely dissolved.

**Figure 4.**
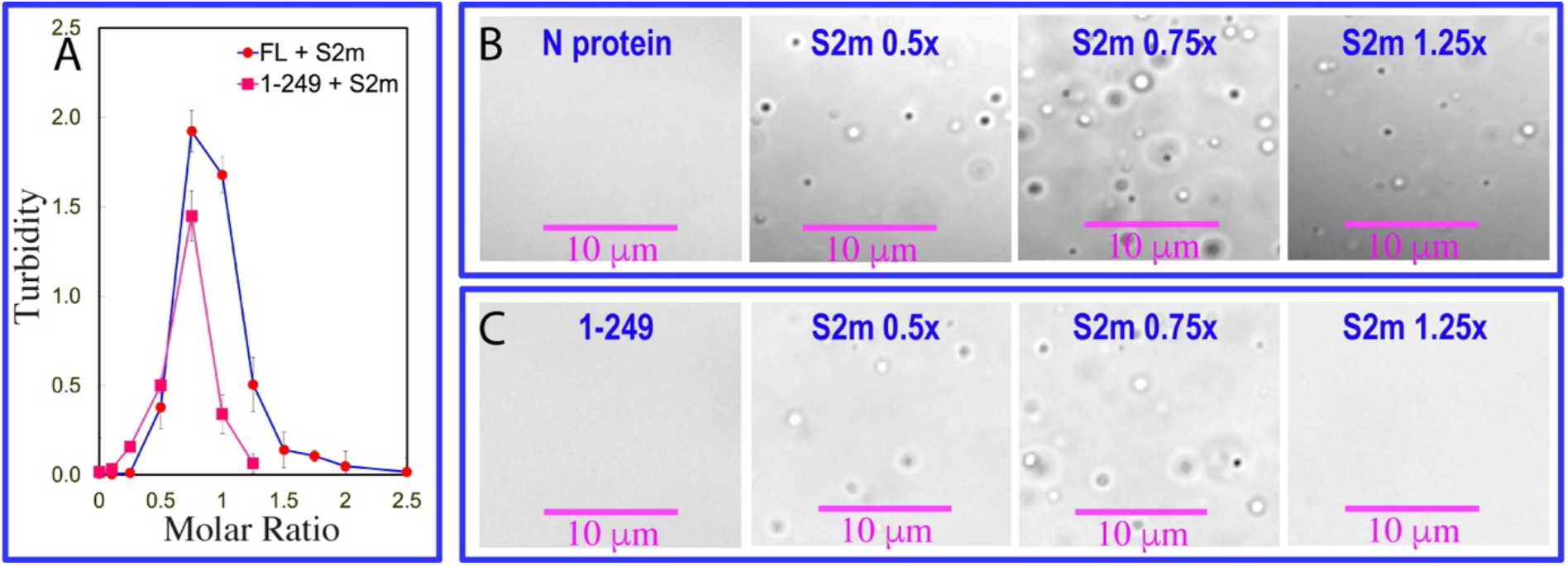
S2m biphasically modulates LLPS of the full-length N and N (1-249) proteins. (A) Turbidity (absorption at 600 nm) curves of the full-length N and N (1-249) proteins in the presence of S2m at different ratios. (B) DIC images of the full-length N protein in the presence of S2m at different ratios. (C) DIC images of N (1-249) protein in the presence of S2m at different ratios.

Subsequently, we titrated S2m into the N (1-249) protein under the same conditions. Again N (1-249) showed no phase separation in the free state. Nevertheless, S2m could also biphasically modulate LLPS of the N (1-249) protein. Briefly, LLPS could be induced upon addition of S2m and at 1:0.75 (1-249:S2m), the turbidity reached the highest of 1.45 (Fig. 4A) and many liquid droplets with the diameter of ∼1 μm were also formed (Fig. 4C). Similar to what was observed above on the full-length N protein, further addition of S2m led to the reduction of turbidity and dissolution of the droplets. At 1:1.25 all liquid droplets were dissolved. Compared with the liquid droplets formed by the full-length N protein in the presence of S2m at 1:0.75, the droplets formed by the N (1-249) protein with S2m at 1:0.75 have similar sizes but the number were less, thus resulting in the lower turbidity (Fig. 4A).

### Residue-specific NMR view of LLPS of N (1-249) modulated by S2m

So far, no high-resolution mechanism has been reported on LLPS of N protein induced by nucleic acid. Here, our successful identification of N (1-249) offered us the unprecedented opportunity to gain residue-specific view of LLPS by NMR spectroscopy. To achieve this, we monitored the S2m-modulated LLPS by NMR upon stepwise addition of S2m into the N (1-249) sample at ratios 1:0.05, 1:0.1, 1:0.25, 1:0.75, 1:1, and 1:2,5, which range from the induction to complete dissolution of liquid droplets.

As shown in I of Fig. 5, even upon addition of S2m at 1:0.05, some HSQC peaks became broadened and consequently their intensity reduced. At 1:0.1 (II of Fig. 5) further broadening was observed for many HSQC peaks. Noticeably, at 1:0.25, HSQC peaks of all folded NTD and some IDR residues became disappeared while at 1:0.75, even the intensity of the remaining peaks of IDR residues became largely reduced (III of Fig. 5). On the other hand however, further addition of S2m to 1:1 led to the intensity increase for remaining IDR HSQC peaks, and at 1:2.5, the intensity of those IDR HSQC peaks further increased (V of Fig. 5). Nevertheless, even at 1:2.5 where liquid droplets were completely dissolved, the HSQC peaks of NTD and some IDR residues still remained undetectable (VI of Fig. 5).

**Figure 5.**
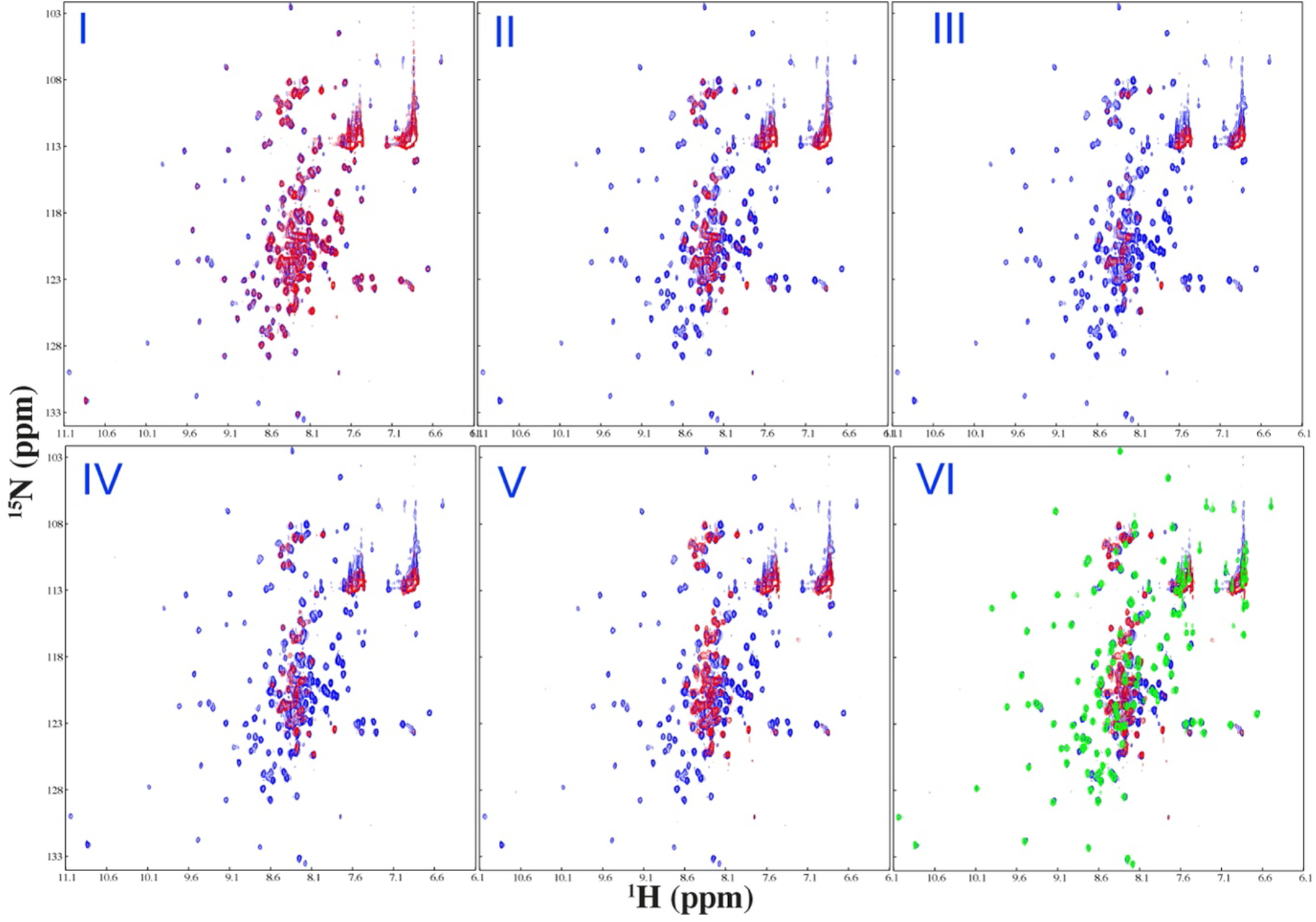
NMR view of the biphasic modulation of LLPS of N (1-249) by S2m. Superimposition of HSQC spectra of N (1-249) in the free state (blue) and in the presence of S2m (red) at 1:0.05 (I), 1:0.25 (II) and 1:0.75 (III), 1:1 (IV) and 1:2.5 (V) (1-249:S2m). (VI) Superimposition of HSQC spectra of N (1-249) in the free state (blue) and in the presence of S2m at 1:2.5 (red) as well as NTD in the free state (green).

Detailed analysis of the peak intensity of the spectra at different S2m ratios reveals a very interesting picture (Fig. 6): as illustrated by Fig. 6A, even with the addition of S2m at 1:0.05, the intensity of the NTD peaks largely reduced with an average of 0.51 while the intensity of IDR peaks showed the relatively small reduction with an average of 0.79. At 1:0.1, the intensity of the NTD peaks further reduced with an average of 0.31 while the intensity of IDR peaks still has an average of 0.73. Strikingly, at 1:0.25, all NTD peaks became too weak to be detected but IDR peaks still has an average of 0.54. At 1:0.75, the intensity of the remaining IDR peaks became further reduced to 0.34. Intriguingly, however, with further addition of S2m to 1:1, the intensity of IDR residues became increased with the average of 0.48 and further back to 0.76 at 1:2.5.

**Figure 6.**
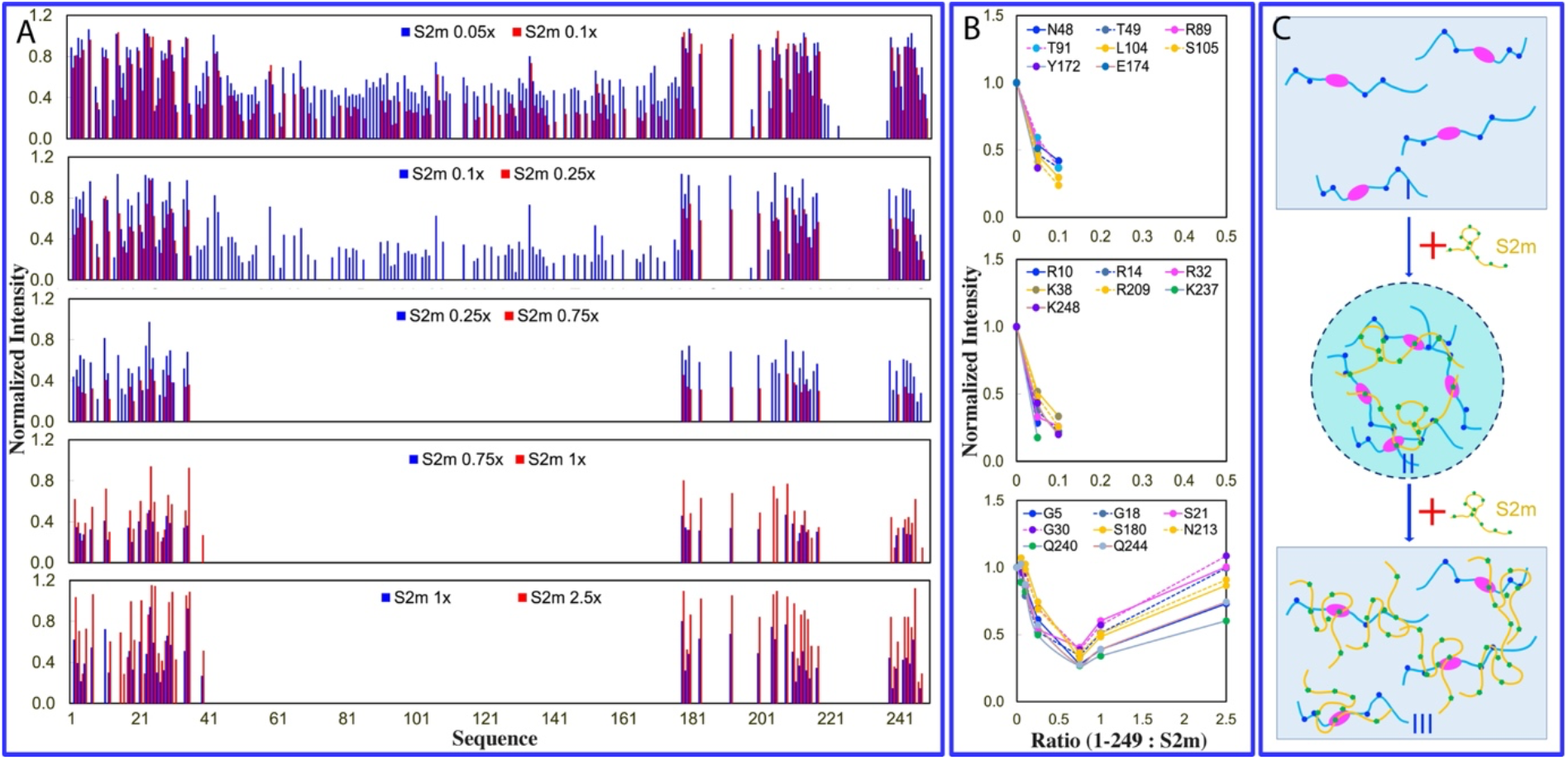
Residue-specific view of the biphasic modulation of LLPS of N (1-249) by S2m. (A) Normalized intensity of N (1-249) residues in the presence of S2m at different ratios as divided by that of N (1-249) residues without S2m. (B) Normalized intensity of three groups of N (1-249) residues in the presence of S2m at different ratios. (C) Speculative model of homogeneous solution of N (1-249) (I) which is induced to phase separate upon adding S2m at low ratios (II) followed by dissolution into homogeneous solution at high ratios (III).

Furthermore, a close examination indicates that based on the patterns of the intensity changes, the N (1-249) residues can be grouped into three categories (Fig. 6B): the folded NTD residues (group 1) and 7 Arg/Lys residues within IDRs (group 2) with their HSQC peak intensity reduced rapidly upon adding S2m and becoming too weak to be detected above 1:0.25, Interestingly those two categories of residues had their HSQC peaks remaining undetectable even with further addition of S2m up to 1:2.5. By contrast, for other IDR residues (group 3), their intensity reduced with addition of S2m and reached the lowest at 1:0.75. However, with further addition of S2m, their peak intensity increased and at 1:2.5, the average intensity is back to 0.76.

In titrations with S2m, the intensity of HSQC peaks of N (1-249) appears to be mechanistically modulated by at least two processes: 1) the formation of large and dynamically crosslinked complexes between S2m and N (1-249) molecules, which is expected to slow the rotational motions and consequently lead to significant broadening of HSQC peaks; and 2) the binding-induced dynamics on μs-ms time scale that also result in significant broadening of HSQC peaks (6,20,24,26,34,35). As NTD is a folded domain, its residues largely behave as a coupled unit. By contrast, due to the lack of the folded structure, IDR residues behave rather independently. As such, upon binding with S2m, HSQC peaks of all NTD residues become uniformly broadened and disappeared due to the μM Kd or/and provoked μs-ms dynamics. By contrast, IDR residues have rather independent dynamic behaviors and therefore only the peaks of Arg/Lys residues directly bound with S2m become significantly broadened and disappeared, while the peaks of other IDR residues only have reduced intensity due to the slowing of the rotational motions upon forming the large dynamically-crosslinked complexes. However, upon further addition of S2m, the large complexes are disrupted and consequently the intensity of most IDR residues except Arg/Lys become increased. Nevertheless, the peaks of NTD and Arg/Lys peaks remain undetectable even in the exceeding presence of S2m because these residues are still bound with S2m even after the complete dissolution of liquid droplets in the presence of the exceeding amount of S2m.

Previously RNA (36) and ssDNA (37) were shown to biphasically modulate LLPS of FUS and TDP-43 respectively but the general mechanisms remain largely elusive. Here the results indicate that S2m appears to achieve both induction and dissolution of LLPS of N (1-249) mainly by specifically binding to the folded NTD and Arg/Lys residues within IDRs. In this context, a speculative model was proposed (Fig. 6C). Briefly, upon adding S2m into the homogenous solution of N (1-249) (I of Fig. 6C) at low ratios, N (1-249) and S2m are capable of dynamically but specifically interacting with each other multivalently over both NTD and IDR Arg/Lys residues to form large and dynamically-crosslinked complexes which manifest as liquid droplets (II of Fig. 6C). However, with the exceeding addition of S2m, several S2m molecules become bound with one N (1-249) molecule and consequently the large and dynamically-crosslinked complexes become disrupted, thus manifesting as the dissolution of liquid droplets into homogeneous solution (III of Fig. 6C).

### ATP dissolves LLPS of the full-length N and N (1-249) proteins in the same manner

The residue-specific NMR results together imply that ATP and S2m might in fact have the highly overlapped binding sites on N (1-249), namely NTD and Arg/Lys of IDRs despite having very different affinities. If this is the case, ATP at the much higher concentrations than that of S2m should be able to dissolve the S2m-induced LLPS of N (1-249) by competitively displacing S2m from being bound with the proteins. To verify this mechanism, we prepared the phase separated samples of both full-length N and N (1-249) proteins with the pre-presence of S2m at 1:0.75. Subsequently, ATP was added into the samples in a stepwise manner, as monitored by turbidity (Fig. 7A) and DIC imaging (Fig. 7B and S5).

**Figure 7.**
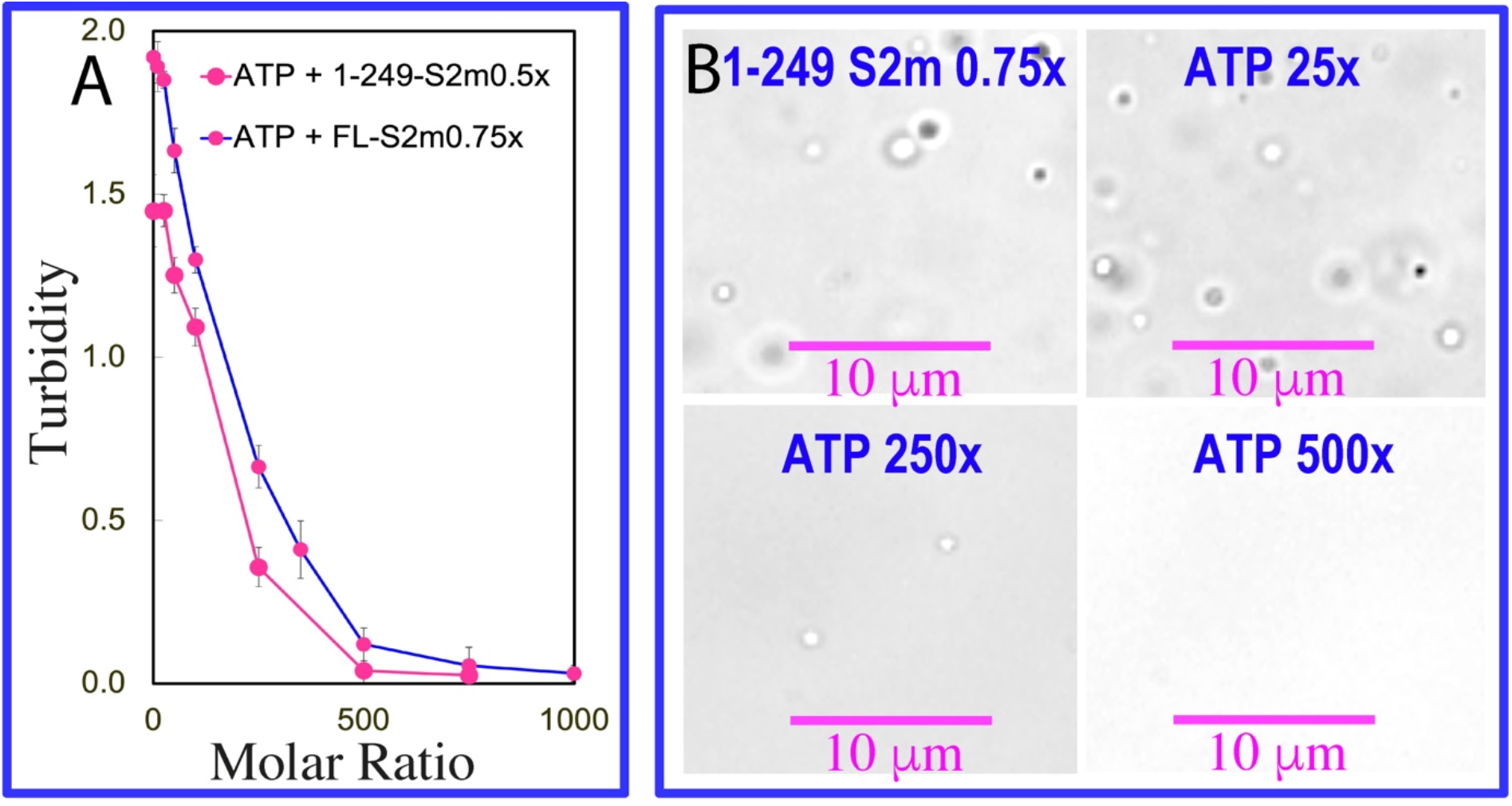
ATP dissolves LLPS of N (1-249) induced by S2m. (A) Turbidity curves of the full-length N protein and N (1-249) in the presence of S2m at 1:0.75 with additional addition of ATP at different ratios. (B) DIC images of N (1-249) in the presence of S2m at 1:0.75 with additional addition of ATP at different ratios.

Indeed, ATP could dissolve LLPS of both full-length N and N (1-249) proteins induced by S2m, although ATP completely dissolved LLPS of N (1-249) at the slightly lower ratio (1:500) than that for dissolving LLPS of the full length N protein (1:750). This difference is most likely due to the presence of CTD in the full-length N protein which might enhance LLPS by additional involvement in binding S2m or/and dimerization/oligomerization. Briefly, as for N (1-249), at the ratio <1:25, ATP only has minor effect on LLPS. However, at ratio of 1:250, ATP significantly dissolved liquid droplets as evidenced by the significant reduction of turbidity and disappearance of many liquid droplets. Strikingly, at 1:500, ATP could completely dissolve liquid droplets of N (1-249) protein induced by S2m. The result clearly suggest that ATP and S2m do interplay in modulating LLPS of the full-length N and N (1-249) proteins in the same manner.

### ATP dissolves LLPS by specifically competing with S2m for binding the protein

To understand the high-resolution mechanism, we monitored the dissolution of the S2m-induced LLPS of N (1-249) by ATP with HSQC spectroscopy. As shown in I of Fig. 8A, the addition of ATP at 1 mM into the N (1-249) sample with the pre-existence of S2m at 1:0.75 has very minor effect on its HSQC spectrum. However, addition of ATP at 10 mM led to the restore of some HSQC peaks (II of Fig. 8A) and at 20 mM, the disappeared HSQC peaks including those of the NTD and Arg/Lys residues were re-appeared (III of Fig. 8A). Most strikingly, the HSQC spectrum in the presence of S2m at 1:0.75 with additional addition of ATP at 20 mM is highly similar to that without S2m but only in the presence of ATP at 10 mM. This observation unambiguously suggests that ATP is able to bind NTD and IDR Arg residues as well as to significantly displace S2m from being bound with the protein. Nevertheless, even at 20 mM, many HSQC peaks, particularly from NTD residues are still weak as compared with those of N (1-249) only in the presence of ATP at 10 mM, implying that ATP even at 20 mM is still unable to completely displace the binding of S2m from N (1-249) because the binding affinity of S2m to N (1-249) is much higher (with Kd of ∼μM) than that of ATP (Kd of ∼mM). However, further addition of ATP to high concentrations led to a significant increase of solution viscosity and thus the universally weakening of NMR signals.

**Figure 8.**
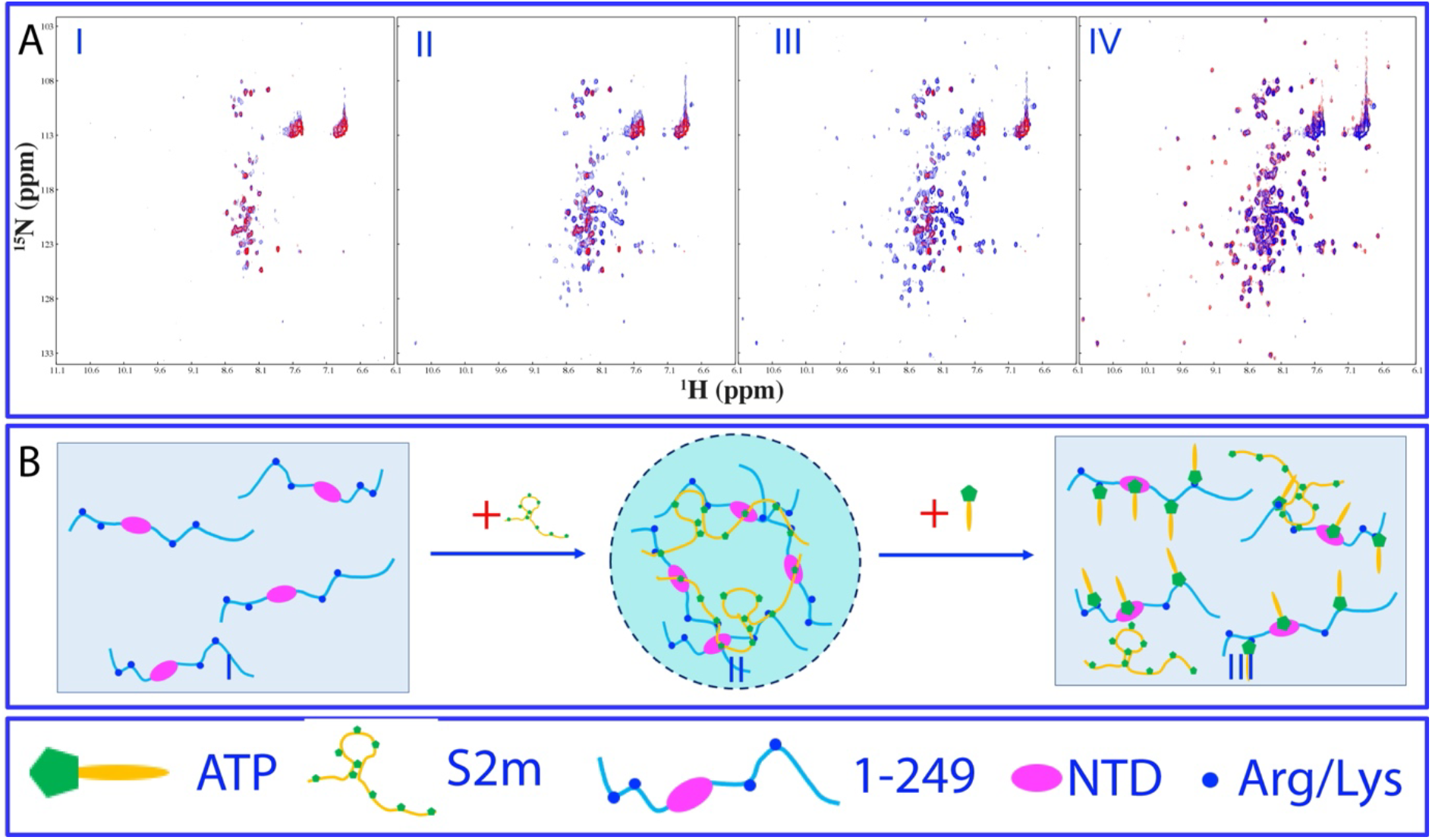
NMR view of the interplay of ATP and S2m in modulating LLPS of N (1-249). Superimposition of HSQC spectra of N (1-249) in the presence of S2m at 1:0.75 (red) with additional addition of ATP (blue) at 5 mM (I), 10 mM (II) and 20 mM (III). (IV) Superimposition of HSQC spectra of N (1-249) in the presence of ATP at 10 mM only (red) and in the presence of both S2m at 1:0.75 and ATP at 20 mM (blue). (B) Speculative model of N (1-249) in homogeneous solution (I) which undergoes phase separation to form dynamic liquid droplets upon induction by S2m (II), followed by dissolution of droplets into homogeneous solution upon adding the exceeding amount of ATP (III).

Here the NMR competition experiments confirm the above speculation that ATP and S2m share the highly-overlapped binding sites over both folded NTD and IDRs of N (1-249). As a consequence, S2m specifically interacts with N (1-249) (I of Fig. 8B) to induce LLPS by forming the large and dynamically crosslinked complexes (II of Fig. 8B), which is characterized by an extensive intensity reduction and disappearance of HSQC peaks. However, the addition of ATP at the high concentrations is able to specifically displace S2m from being bound with N (1-249), thus leading to the dissolution of the liquid droplets (III of Fig. 8B), with the reappearance of HSQC peaks including those of the NTD and Arg/Lys residues. Nevertheless, due to the fact that S2m binds N (1-249) with Kd of ∼μM, which is much higher that that of ATP (Kd of ∼mM), ATP at 20 mM is still unable to completely displace S2m from binding N (1-249), and consequently the intensity of HSQC peaks of N (1-249) in the presence of both S2m and ATP is still weaker than that only in the presence of ATP.

### S2m modulates LLPS of N (175-419) protein

We also assess whether S2m could induce LLPS for N (175-419), N (175-365) and N (247-419). As shown in Fig. 9A, the N (175-419) sample showed no LLPS in the free state but phase separation was induced upon addition of S2m as evident by the increase of turbidity and DIC imaging. At 1:0.5, the turbidity reached the highest of 1.71 (I of Fig. 9A) and many liquid droplets with the diameter of ∼1 μm were formed (II of Fig. 9A). However, as observed on N protein and N (1-249) above, further addition of S2m led to the reduction of turbidity and dissolution of the droplets. At 1:1.25 all liquid droplets were completely dissolved.

**Figure 9.**
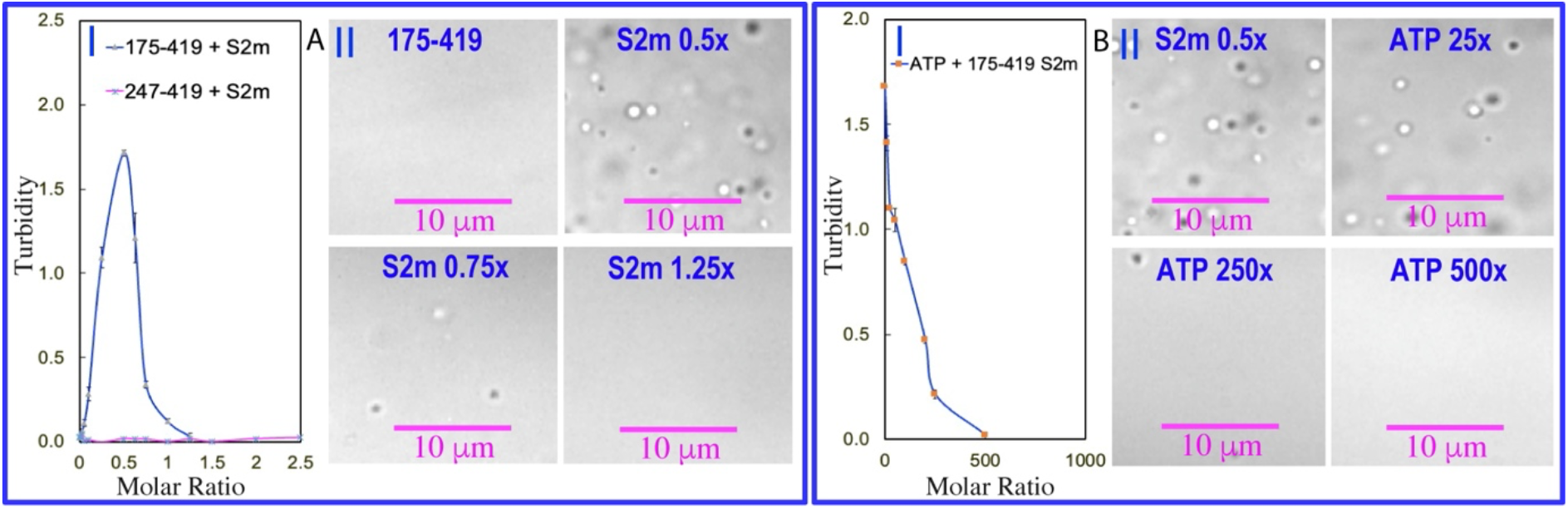
S2m and ATP modulate LLPS of N (175-419). (A) Turbidity curves of N (175-419) in the presence of S2m at different ratios (I); and DIC images of N (175-419) in the presence of S2m at different ratios. (B) Turbidity curves of N (175-419) in the presence of S2m at 1:0.5 with additional addition of ATP at different ratios (I); and DIC images of N (175-419) in the presence of S2m at 1:0.5 with additional addition of ATP at different ratios (II).

On the other hand, for N (175-365), addition of S2m even at 1:0.1 led to visible precipitation and consequently no DIC characterization could be performed. For N (247-419), addition of S2m even up to 1:2.5 induced no LLPS as well as precipitation as reported by turbidity (I of Fig. 9A). The results suggest that CTD needs the additional presence of IDR2 with 8 Arg and 3 Lys residues to achieve multivalent and dynamic binding of S2m with CTD and Arg/Lys, which is sufficient to drive LLPS. By contrast, as IDR3 contains only 1 Arg and 9 Lys, the interaction of S2m with Arg/Lys is not sufficient for driving LLPS as the interaction between nucleic acids and Lys is much weaker that that with Arg (23,38).

We also prepared the phase separated samples of N (175-419) with the pre-presence of S2m at 1:0.5. Subsequently, ATP was added into the samples in a stepwise manner, as monitored by turbidity (I of Fig. 9B) and DIC imaging (II of Fig. 9B). Indeed, ATP could dissolve its LLPS induced by S2m and the droplets were completely dissolved at 1:500, which is in general similar to what was observed on N (1-249) and full-length N protein.

### NMR characterization of the interaction of N (247-419) and S2m

We further set to characterize the interaction of S2m with N (175-419), N (175-365) and N (247-419) by NMR. Unfortunately, as to collect good quality NMR spectra needs a protein concentration of at least 50 μM, we were unable to collect NMR spectra of (175-419) and N (175-365) as they became precipitated at 50 μM upon adding S2m even at 1:0.1. Nevertheless, we have successfully collected NMR spectra of N (247-419) at 50 μM in the presence of S2m up to 1:2.5 (SFig. 6). Intriguingly, although the addition of S2m triggered no LLPS even at 50 μM, S2m was shown by NMR (Fig. S6) to interact with the folded CTD as evidenced by the broadening of several very up-field NMR signals, exactly as we previously observed on the interaction of S2m with the isolated CTD (27). On the other hand, as judged by its HSQC spectra in the presence of S2m at different concentrations, most HSQC peaks of the IDR3 residues in N (247-419) retained even up to 1:2.5, implying that the interaction of S2m with IDR3 is indeed relatively weak as compared to that between S2m and Arg-rich IDR1 and IDR2 which resulted in dramatic disappearance of many of their HSQC peaks (Fig. 5 and Fig. 6).

## Discussion

Recently accumulated results suggest that LLPS is also emerging as the common mechanism underlying key steps of the life cycle of viruses more than SARS-CoV-2, which include HIV, Measles and Hendra viruses (36-40). Intriguingly, viral proteins are rich in IDRs which have been extensively identified to phase separate upon interacting with the viral or/and host RNA/DNA (39-43). Therefore the delineation of the critical roles of LLPS not only sheds light on previously-unknown principles underlying the host-virus interaction and viral life cycle, but most importantly opens up a new direction for development of antiviral strategies/drugs although this still remains almost completely unexplored so far. Nevertheless, this might be a very challenging task because it requires the understanding of the mechanisms of the viral LLPS at high resolution which is, however, characterized by two unusual properties: 1) the proteins undergoing LLPS are rich in IDRs; 2) the protein-nucleic-acid complexes manifesting as liquid droplets are multivalent and highly dynamic. In this context, the molecular mechanisms of LLPS become not amenable for high-resolution investigations by many classic biophysical methods such as X-ray crystallography.

More specifically, three key issues need to be addressed in order to build up a basic framework for understanding LLPS of SARS-CoV-2 N protein: 1) what is the exact role of IDRs in driving LLPS? 2) is the interaction between proteins and nucleic acids highly specific or just non-specific salt/buffering effects? 3) is there any host-cell small molecule, which can specifically modulate the viral LLPS? In the present study, we aimed to address them associated with LLPS of SARS-CoV-2 N protein by NMR, the only biophysical method so far which can provide residue-specific knowledge of LLPS, in aid with turbidity measurements and DIC imaging.

Extensive studies, for the first time, to the best of our knowledge, decode that S2m, a SARS-CoV-2-specific nucleic acid probe, induces and then dissolves LLPS of N protein mainly by dynamic but specific interactions multivalently over both folded NTD/CTD and Arg/Lys residues within IDRs. In particular, our studies provide the first residue-specific evidence that Arg/Lys residues within IDRs are also capable of directly binding with nucleic acids. As such, in addition to passively providing the essential dynamics for forming liquid-like droplets, IDRs in fact directly drive LLPS by utilizing their Arg/Lys residues to dynamically interact with nucleic acids. Therefore, LLPS of SARS-CoV-2 N protein is mainly driven by the specific interactions of nucleic acids dynamically and multivalently with the protein. However, nucleic acids regardless of types and sequences appear to be sufficient to establish the interactions for driving LLPS with proteins containing the folded nucleic-acid-binding domains NTD/CTD or/and Arg residues within IDRs. The presence of CTD and IDR2 led to proneness to oligomerization in N (175-419) and the full-length N protein and consequently might change the strength of the driving force or/and material properties of liquid droplets. In the general sense, the molecular mechanisms for ATP and nucleic acids to modulate LLPS of SARS-CoV-2 N and human FUS proteins (21,22), both of which contain the nucleic-acid-binding domains and Arg/Lys-rich IDRs are highly conserved although their biological roles are fundamentally distinctive.

Previously RG-/RGG-motifs within IDRs have been identified to bind nucleic acids without a strict requirement of types and sequences (20,44-46). Here our NMR results showed that none of four IDR Arg residues of N protein interacting with nucleic acid are within the RG-/RGG-motifs but instead with very diverse sequences: Gln9-Arg10-Asn11; Pro13-Arg14-Ile15; Glu31-Arg32-Ser33; and Ala208-Arg209-Met210. Therefore our results extend the previous notion by showing that as long as Arg residue is located within IDRs and accessible, it is capable of interacting nucleic acids without needing to be within RG-/RGG motifs. On the other hand, Lys38, Lys237 and Lys248 were also identified to interact with nucleic acid, in general consistent with the previous report that Lys residue is critical for interacting with RNA to drive LLPS of the human proteins in P-body (38). The binding affinity of nucleic acids to Arg appears to be higher than that to Lys because the aromatic base rings of nucleic acids can only establish Τ-cation interaction with the side chain of Lys residue, but Τ-Τ and Τ-cation interactions with the side chain of Arg residue (20,44). As such, N (247-419) with 6 Lys but only 1 Arg in IDR3 has no capacity in phase separation although S2m is still able to bind the folded CTD.

Most unexpectedly, ATP with concentrations > mM in all living cells but totally absent in viruses, has been previously decoded to bind both the folded NTD and CTD with the Kd of 3.3 ± 0.4 and 1.49 ± 0.28 mM respectively. Here we showed that in the context of N (1-249), ATP binds NTD with a slightly high affinity with Kd of 2.0 ± 0.2 mM. Most importantly, for the first time we revealed that Arg residues within IDRs of a viral protein have the comparable affinity with Kd of 2.8 ± 0.2 mM). Most critically, ATP acts to dissolve the S2m-induced liquid droplets by specifically displacing S2m from being bound with the protein to disrupt the large multivalently and dynamically-crosslinked S2m-protein complex, thus clearly indicating that ATP and S2m share the highly overlapped sites over both folded NTD and IDRs of the viral N protein, in generally similar to what we previously observed on human FUS protein (20,22). Nevertheless, the much higher concentration of ATP is needed to dissolve the S2m-induced LLPS of N protein, because unlike ATP with only one aromatic base ring and one triphosphate group, the nucleic acid fragment contains many flexibly-linked aromatic base rings and phosphate groups. Consequently one nucleic acid fragment is sufficient to establish the multivalent binding with the folded NTD or/and IDRs of N protein, thus leading to the much higher affinity (47). Nevertheless, the specific capacity of ATP in dissolving the S2m-induced LLPS of N protein bears significant biological relevance because in the host cell ATP always has >mM concentrations but at the initial stage of infection, the number of viral gRNA and N protein should be very low based on the estimation that in one SARS-CoV-2 virion, ∼730– 2200 copies of N protein form the complex with one 30-kb gRNA (48). As such, upon immediate entry into the host cell of the virus, ATP is expected to facilitate the uncoating of the SARS-CoV-2 gRNA-N-protein condensates which is essential for initiating the viral life cycle in the host cell. In this sense, ATP appears to be evolutionarily hijacked by SARS-CoV-2 to promote its own life cycle in the host cell and thus emerges a key cellular small molecule which critically controls the host-SARS-CoV-2 interaction.

Pandemic-relevantly, the sequences of SARS-CoV-2 N protein of different variants are highly conserved not only over the folded NTD and CTD, but also over Arg/Lys residues within IDRs. This implies that the mechanisms underlying LLPS of N protein as specifically induced by nucleic acids and dissolved by ATP might be highly conserved in different variants including Delta and Omicron. Therefore ATP could serve as a key lead for further design of anti-SARS-CoV-2 drugs efficient for different variants by disrupting the highly conserved LLPS of N protein. Interestingly, Imatinib, one of three drug candidates WHO was set for clinical trial to treat SARS-CoV-2, is an ATP mimetic originally used to treat cancer by inhibiting tyrosine kinases (49). However, its mechanism to inhibit SARS-CoV-2 remains completely unknown and therefore it would be of significant interest to investigate whether it can also modulate LLPS of SARS-CoV-2 N protein. As the properties of SARS-CoV-2 N protein required for interacting with nucleic acids to drive LLPS can be extensively found in the IDR-rich proteins of other viruses, it is thus tempting to speculate the mechanisms of LLPS deciphered here as specifically induced by nucleic acids and dissolved by ATP might be also conserved to a certain degree in other viruses. If this is confirmed, it is possible that ATP and ATP-like small molecules might have the common ability to modulate the viral LLPS essential for their life cycle and most important could serve as leads for design of antiviral drugs for other viruses.

In conclusion, we have successfully identified N (1-249) to be amenable for high-resolution NMR investigations and this allowed us to gain residue-specific insights into the mechanism of LLPS of SARS-CoV-2 N protein which is specifically induced by nucleic acids and dissolved by ATP. The results decipher that LLPS of N protein is specifically driven by dynamically and multivalently interactions of nucleic acids with the folded nucleic-acid-binding NTD/CTD and Arg/Lys residues within IDRs. Furthermore, ATP acts to dissolve LLPS by specifically competing with nucleic acids for binding the highly overlapped sites of the protein, thus disrupting the dynamic complex. Most strikingly, as the residues critical for binding nucleic acids and ATP are highly conserved not only over the folded NTD/CTD, but also over Arg/Lys residues within IDRs in different SARS-CoV-2 variants. ATP might also serve as a promising lead to develop small molecule drugs efficient for different variants by disrupting the highly conserved LLPS of SARS-CoV-2 N protein. Furthermore, because LLPS of IDR-containing viral proteins induced by nucleic acids is increasingly identified in other viruses, in the future it is of extreme significance to explore whether the mechanisms we decoded here are also conserved in other viruses. In particular, as ATP universally has very high concentrations in all living cells but is absent in all viruses, it is of significant interest to explore whether ATP might be a universal endogenous small molecule to control the host-virus interactions. In the general framework, the mechanisms of LLPS modulated by ATP and nucleic acids appear to be conserved from human to viral proteins, which are all dependent on multivalent dynamic but specific interactions of nucleic acids with the folded nucleic-acid-binding domains and Arg/Lys within IDRs.

## Materials and methods

### Preparation of recombinant SARS-CoV-2 N protein as well as its dissected domains

The gene encoding 419-residue SARS-CoV-2 N protein was purchased from a local company (Bio Basic Asia Pacific Pte Ltd), which was cloned into an expression vector pET-28a with a TEV protease cleavage site between N protein and N-terminal 6xHis-SUMO tag used to enhance the solubility (10). The DNA fragments encoding its N (1-249), N (175-419), N (175-364) and N (247-419) as well as NTD (44-180) and CTD (247-364) were subsequently generated by PCR rection and subcloned into the same vector.

The recombinant full-length N protein and its dissected domains were expression in *E. coli* cells BL21 with IPTG induction at 18 ºC, which were found to be soluble in the supernatant. For NMR studies, the bacteria were grown in M9 medium with addition of (^15^NH_4_)_2_SO_4_ for _15_N-labeling. The recombinant proteins were first purified by Ni^2+^-affinity column (Novagen) under native conditions and subsequently in-gel cleavage by TEV protease was conducted. The eluted fractions containing the recombinant proteins were further purified by FPLC chromatography system with a Superdex-200 column for the full-length, N (1-249), N (175-419), N (175-364) and N (247-419) as well as a Superdex-75 column for NTD and CTD (50). The purity of the recombinant proteins was checked by SDS-PAGE gels and NMR assignments (51,52). ATP was purchased from Merck. Protein concentration was determined by spectroscopic method in the presence of 8 M urea (53).

### NMR characterizations of differentially-dissected domains

Due to the proneness of self-association, after extensive screening of protein concentrations and buffer conditions, the samples of the full-length N and N (175-419) proteins were prepared in 25 mM HEPES buffer (pH 6.5) with 70 mM KCl, while the samples of the other constructs were in 10 mM sodium phosphate buffer (pH 6.5) in the presence of 150 mM NaCl, which mimics the cellular environments. NMR experiments were conducted at 25 ºC on an 800 MHz Bruker Avance spectrometer as described previously (10,20,21,27,54).

### NMR characterizations of the binding of ATP and S2m to N (1-249) and NTD

NMR samples of N (1-249), N (175-419), and NTD were prepared at 100 μM in 10 mM sodium phosphate buffer (pH 6.5) in the presence of 150 mM NaCl. ATP was dissolved in the same buffer (10,18,20,21,27). The final solution pH was adjusted to 6.5 by use of very diluted HCl or NaOH.

For NMR titrations to determine residue-specific Kd of N (1-249) residues for binding ATP, two dimensional ^1^H-^15^N NMR HSQC spectra were collected on the ^15^N-labelled sample at 100 μM in the presence of ATP at 0, 0.5, 1, 2, 4, 6, 8, 10 mM. For NMR characterization of the binding of N (1-249) with S2m, HSQC spectra were collected on the ^15^N-labelled sample at 100 μM with addition of S2m at 0, 1:0.05, 1:0.1, 1:0.25, 1:0.75, 1:1, 1:2.5 (1-249:S2m). For NMR characterization of the binding of N (175-419), N (175-364) and N (247-419) with S2m, HSQC spectra were collected on the ^15^N-labelled sample at 50 μM. For NMR investigation on the interplay of ATP and S2m in modulating LLPS of N (1-249), HSQC spectra were collected on the phase separated sample of the ^15^N-labelled N (1-249) at 100 μM in the pre-existence of S2m at 1:0.75, into which ATP was subsequently added at different combinations. NMR data were processed with NMRPipe (55) and analyzed with NMRView (56).

### Calculation of CSD and data fitting

Sequential assignments were achieved based on the deposited NMR chemical shifts for NTD and N (1-249) (30,31). To calculate chemical shift difference (CSD), HSQC spectra collected without and with ATP or S2m at different concentrations were superimposed. Subsequently, the shifted HSQC peaks were identified and further assigned to the corresponding residues. The chemical shift difference (CSD) was calculated by an integrated index with the following formula (10,18,20,21,27,34):

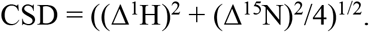

In order to obtain residue-specific dissociation constant (Kd), we fitted the shift tracings with significant shifts (CSD > average + STD) by using the following formula (10,21,34):

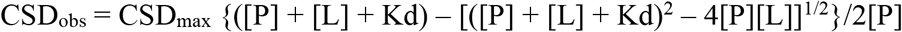

Here, [P] and [L] are molar concentrations of CTD and ligands (ATP) respectively.

### LLPS imaged by differential interference contrast (DIC) microscopy

The formation of liquid droplets was imaged on 50 μl of the full-length N protein or N (1-249) samples by DIC microscopy (OLYMPUS IX73 Inverted Microscope System with OLYMPUS DP74 Color Camera) as previously described (10,20,27). The protein samples were prepared at 20 μM in 25 mM HEPES buffer (pH 6.5) with 70 mM KCl. The turbidity (absorption at 600 nm) were measured for all DIC samples with three repeats.

## Acknowledgement

This study is supported by Ministry of Education of Singapore (MOE) Tier 1 Grants R-154-000-B92-114 to Jianxing Song.

## Author contributions

Conceived the research: J.S. Performed research and analyzed data: M.D, T.L. and J.S; Acquired funding: J.S; Wrote manuscript: J.S.

